# Topography of inputs into the hippocampal formation of a food-caching bird

**DOI:** 10.1101/2023.03.14.532572

**Authors:** Marissa C. Applegate, Konstantin S. Gutnichenko, Dmitriy Aronov

**Affiliations:** Zuckerman Mind Brain Behavior Institute, Columbia University

## Abstract

The mammalian hippocampal formation (HF) is organized into domains associated with different functions. These differences are driven in part by the pattern of input along the hippocampal long axis, such as visual input to the septal hippocampus and amygdalar input to temporal hippocampus. HF is also organized along the transverse axis, with different patterns of neural activity in the hippocampus and the entorhinal cortex. In some birds, a similar organization has been observed along both of these axes. However, it is not known what role inputs play in this organization. We used retrograde tracing to map inputs into HF of a food-caching bird, the black-capped chickadee. We first compared two locations along the transverse axis: the hippocampus and the dorsolateral hippocampal area (DL), which is analogous to the entorhinal cortex. We found that pallial regions predominantly targeted DL, while some subcortical regions like the lateral hypothalamus (LHy) preferentially targeted the hippocampus. We then examined the hippocampal long axis and found that almost all inputs were topographic along this direction. For example, the anterior hippocampus was preferentially innervated by thalamic regions, while posterior hippocampus received more amygdalar input. Some of the topographies we found bear resemblance to those described in the mammalian brain, revealing a remarkable anatomical similarity of phylogenetically distant animals. More generally, our work establishes the pattern of inputs to HF in chickadees. Some of these patterns may be unique to chickadees, laying the groundwork for studying the anatomical basis of these birds*’* exceptional hippocampal memory.

## Introduction

The hippocampus is necessary for memory formation in species from every class of vertebrates (Scoville and Milner, 1957; Krushinskaya, 1966; Rodríguez et al., 2002; Inés Sotelo et al., 2022). Different types of memory rely on the hippocampus. For example, rodents need their hippocampus for both *‘*spatial*’* memory (e.g., location of a hidden platform in the Morris water maze) and *‘*emotional*’* memory (e.g., link of a tone with a foot shock) (Morris et al., 1982; Kjelstrup et al., 2002; Maren and Holt, 2004). These distinct functions are anatomically organized along the dorso-ventral (DV) *“*long*”* axis of the hippocampus (Moser and Moser, 1998; Fanselow and Dong, 2010; Strange et al., 2014). The dorsal hippocampus is needed for spatial memory tasks (Cimadevilla et al., 2000; Barbosa et al., 2012), and neural recordings there have shown spatially tuned cells (Jung et al., 1994; Kjelstrup et al., 2008). In contrast, the ventral hippocampus has been implicated in a variety of emotional tasks, and neurons there exhibit far less spatial tuning (Henke, 1990; Kjelstrup et al., 2002).

In non-mammalian species, the hippocampus is also involved in both spatial and emotional behaviors (Krushinskaya, 1966; Sherry and Vaccarino, 1989; Reis et al., 1999; Damphousse et al., 2022). Anatomical organization of these functions seems to be similar to that in mammals. In birds, the hippocampal long axis extends in an antero-posterior (AP) direction, with anterior hippocampus likely homologous to the rodent dorsal hippocampus (Smulders, 2017; Herold et al., 2019; Payne et al., 2021; Damphousse et al., 2022). Anterior hippocampus contains spatially tuned neurons and is preferentially involved in spatial tasks, whereas posterior hippocampus is less spatially tuned and is involved in stress-related functions.

What drives these functional differences across the hippocampus? In mammals, functional organization is driven in part by the anatomy of inputs. For example, head direction signals from the anterior thalamus preferentially target the dorsal HF, where they likely contribute to spatial representations (Prasad and Chudasama, 2013). The amygdala, which has been implicated in emotionally driven behaviors, innervates ventral hippocampus and likely contributes to the emotionally linked deficits of ventral-hippocampal lesions (Van Groen and Wyss, 1990). However, it is unknown whether a similar organization of inputs along the long axis exists in non-mammals.

Further functional segmentation of the hippocampal formation (HF) occurs along the transverse axis. In mammals, there are major differences in input between the hippocampus and the entorhinal cortex. For example, cortical inputs preferentially innervate the entorhinal cortex (Burwell and Amaral, 1998) while hypothalamic inputs preferentially innervate the hippocampus (Swanson and Cowan, 1975; Hahn and Swanson, 2012). These anatomical differences may have functional consequences, possibly driving differences in spatial representations – like hippocampal place cells and entorhinal grid and border cells (O*’*Keefe and Dostrovsky, 1971; Hafting et al., 2005). Recent work in birds has revealed a similar functional organization along the transverse axis. The avian hippocampus contains abundant place cells, while the dorsolateral hippocampal formation (DL) contains many entorhinal-like firing patterns (Agarwal et al., 2021; Payne et al., 2021; Applegate et al., 2023). Again, it is unknown how the patterns of inputs into these regions compare to mammals.

We decided to study the organization of inputs to HF in a food-caching bird, the black-capped chickadee (*Poecile atricapillus*). These animals are memory specialists, capable of remembering hidden locations of thousands of food items scattered throughout their environment (Pravosudov, 1985; Stevens and Krebs, 1986). These memories require the hippocampus (Krushinskaya, 1966; Sherry and Vaccarino, 1989). Previous studies have examined avian inputs to HF (Kraniak and Siegel, 1978; Casini et al., 1986; Székely and Krebs, 1996; Atoji et al., 2002; Atoji and Wild, 2004), but no anatomical tracing of HF has been done in a food-caching bird. In addition to analyzing inputs along the two axes of HF, we were interested in what may be unique to food-caching species. The hippocampus of food-caching birds is hypertrophic, roughly three times larger than the hippocampus of non-food-caching birds (Krebs et al., 1989; Garamszegi and Eens, 2004). There could be interesting differences in connectivity as well, further motivating the study of inputs to HF in food-caching birds.

### Abbreviations of anatomical terms

Cb: Cerebellum
CDL: Caudal dorsolateral hippocampal formation
DL: Dorsolateral hippocampal formation
DLM: Nucleus dorsolateralis anterior thalami, pars medialis
DTB: Diencephalo-telencephalic boundary
E: Entopallium
FDB: Fascicle of the diagonal band of Broca
FLM: Fasciculus longitudinalis medialis
HD: Hyperpallium densocellulare
HF: Hippocampal formation
Hp: Hippocampus
Hy: Hypothalamus
LC: Nucleus linearis caudalis
LFM: Lamina frontalis suprema
LFS: Lamina frontalis superior
LHy: Lateral hypothalamus
LM: Lamina mesopallialis
LoC: Locus coeruleus
LPS: Lamina pallio-subpallialis
m: Midline
MS: Medial septum
NDB: Nucleus of the diagonal band of Broca
NFL: Fronto-lateral nidopallium
NIII: Oculomotor nerve
NIL: Lateral intermediate nidopallium
NIL_L_: Lateral intermediate nidopallium, lateral portion
NIL_M_: Lateral intermediate nidopallium, medial portion
NL: Lateral nidopallium
OM: Tractus occipitomesencephalicus
PMI: Nucleus paramedianus interni thalami
S: Septum
SPC: Nucleus superficialis parvocellularis
SRt: Nucleus subrotundus
TnA: Nucleus taeniae
V: Ventricle
VTA: Ventral tegmental area

## Methods

### Experimental model and subjects

All animal procedures were approved by the Columbia University Institutional Animal Care and Use Committee and carried out in accordance with the US National Institutes of Health guidelines. The subjects were 22 black-capped chickadees (*Poecile atricapillus*) collected from multiple sites in New York State using Federal and State scientific collection licenses. Subjects were at least 4 months old at the time of the experiment, but age was not determined more precisely.

Chickadees are not visibly sexually dimorphic; therefore, all surgical procedures were performed blindly to sex. Sex was determined post-mortem. Seven female and 14 male birds were used for retrograde tracing, and one male was used to create a 3D rendering of the brain.

Prior to experiments, all birds were housed in an aviary in groups of 1-3, on a *‘*winter*’* light cycle (9 h:15 h light:dark). Birds were given an ad-libitum supply of a small-bird diet (Small Bird Maintenance Diet, 0001452, Mazuri or Organic High Potency Fine Bird foods, HPF, Harrison*’*s Bird Foods). Upon transfer to the lab for experiments, birds had primary flight feathers trimmed and were individually housed.

### Surgery

Surgeries were performed on birds after 2-16 months (9.6 ± 1.0, mean ± SEM) in captivity, using the same procedures as in our prior work (Applegate et al., 2023). For all surgical procedures, birds were anesthetized with 1-2% isoflurane in oxygen. Feathers were removed from the surgical site, and the skin was treated with Betadine. Birds were then placed into a stereotaxic apparatus for the duration of the surgery. The head was rotated to an angle appropriate for each experiment (Table 1), measured as the angle between the groove in the bone at the base of the upper beak mandible and the horizontal table surface. Throughout the surgery, saline was injected subcutaneously to prevent dehydration (∼0.4 mL every 30-60 min). After the surgery, an analgesic (buprenorphine, 0.05 mg/kg) was injected intraperitoneally.

**Table 1:** Summary of CTB injections. Coordinates and the number of annotated injections for all injection locations. Antero-posterior (AP) and medio-lateral (ML) coordinates are relative to lambda. Dorso-ventral (DV) coordinate is the depth relative to the brain surface. The number of pallial and non-pallial annotations is sometimes different due to birds in which the same wavelength of fluorophore was used bilaterally. In this case, only pallial regions were annotated, since they do not project contralaterally.

| Location index | Region | Coordinate (mm) |  |  |  | Number of annotated injections |  |
| --- | --- | --- | --- | --- | --- | --- | --- |
|  |  | AP | ML | DV | Head angle | Pallial | Non-pallial |
| 1 | Hp | -1.0 | 1.9 | 0.4 | 90° | 4 | 3 |
| 2 |  | 1.0 | 0.4 | 0.4 | 75° | 5 | 4 |
| 3 |  | 3.0 | 0.4 | 0.4 | 75° | 7 | 4 |
| 4 |  | 5.0 | 0.4 | 1.0 | 75° | 8 | 3 |
| 5 | DL | 2.0 | 2.3 | 0.4 | 75° | 5 | 4 |
| 6 |  | 3.5 | 1.7 | 0.4 | 75° | 4 | 4 |
| 7 |  | 5.5 | 0.7 | 0.3 | 75° | 4 | 4 |

### 3D brain reconstruction

To visualize the 3D arrangement of brain structures, we constructed a template chickadee brain. In one bird, stiff wire (Malin Co., 0.006*”* music wire) was inserted horizontally from the lateral wall of the brain toward the midline to create two visible tracts. The bird was immediately perfused (see below), and following tissue fixation, the brain was sliced sagittally. Slices were annotated (Adobe Illustrator) and aligned using custom MATLAB code, with the two lesions used for alignment.

### Anatomical tracing experiments

For retrograde tracing, we used cholera toxin subunit B (CTB in 1X PBS, 0.4% weight/volume) fluorescently conjugated to Alexa Fluor in one of three different wavelengths (488 nm, 555 nm, and 647 nm; C34775, C34776, C34778, respectively, Invitrogen). We made injections into 1-4 locations per animal, using different wavelengths and sometimes different hemispheres for different locations in individual birds.

Injection coordinates for different brain regions are given in Table 1. The majority of the injection locations (2-7) were made at a 75° head angle. However, to access the very posterior Hp (location 1), the brain was rotated to a 90° head angle. If multiple injections were performed, lambda was relocated for each injection.

Hp injections (locations 1-4) were chosen to be evenly spaced along the AP axis. DL injections (locations 5-7) were chosen to be roughly at the centers of those portions of DL that were retrogradely labeled by Hp injections at locations 2-4 (Applegate et al., 2023). In other words, locations 2-4 were topographically matched to locations 5-7. For location 1, the corresponding portion of DL was too small for reliable injections and was omitted from this study.

### Histology and histological imaging

All perfusions were performed 7 days after surgery. Animals were administered ketamine and xylazine (10 mg/kg and 4 mg/kg respectively), then transcardially perfused with 1X PBS followed by 4% paraformaldehyde in PBS. The brains were extracted and stored in 4% paraformaldehyde in PBS for 2 days. They were then sectioned coronally into 100 µm-thick slices (Leica VT1000S vibratome). Sections were stained with fluorescent DAPI (300 nM in 1X PBS, D1306, Invitrogen) and mounted in Vectashield mounting medium (H-1400-10, Vector Laboratories). Sections were then imaged using a AZ100 Multizoom Slide Scanner (Nikon), using filters for DAPI and all injected fluorophores.

### Quantification and statistical analysis

#### Quantification of labeled cells in anatomical tracing

To visualize brain areas in 3D stereotaxic coordinates, we needed to align the images of each coronal section to the reference chickadee brain (described above). First, we found the coronal section corresponding to the center of one of the CTB injections. Since injections were at known stereotaxic coordinates, and the coronal sections were each 100 µm thick, we used this reference point to assign an AP position to each section. Next, we manually annotated the dorsal and ventral endpoints of the midline in each section. These were used to rotated the image of each section to make the midline vertical. The midline was assigned to be at 0 mm ML. The hemisphere ipsilateral to the injection was designated as positive ML, and the contralateral hemisphere was designated as negative ML. We then annotated the dorsal brain surface by using the MATLAB functions *imbinarize* and *bwboundaries*. This brain surface was used to manually align the sections dorso-ventrally to the reference brain. Finally, small manual corrections were made dorso-ventrally and medial-laterally to ensure that the reconstructed brain surface was smooth.

Within these aligned images, we detected all retrogradely labelled cells using a standard Difference-of-Gaussians algorithm for blob detection. We used a series of five sigma values, logarithmically spaced from 2.5 to 12.5 µm – corresponding to blob sizes of 5 to 25 µm. The image of a brain section was smoothed with Gaussians of each of these sigma values. Successive pairs of smoothed images were subtracted from one another to compute four difference-of-Gaussians, and these differences were stacked into a 3-dimensional matrix. Local maxima exceeding 0.02 in this matrix were detected as the centers of the blobs. If two detected blobs were overlapping, the one with the lower value was eliminated.

Slices were then annotated manually to assign detected cells to regions of interest. The boundaries of these brain regions were determined by comparing the DAPI-stained images to published atlases from other species (Karten and Hodos, 1967; Stokes et al., 1974; Puelles et al., 2019). We found 15 regions outside of HF in which there was retrograde labeling resulting from every injection in at least one of the coordinates (Table 1). These regions were annotated and analyzed. Labeled cells in other regions were never observed following more than one injection in one bird. These were usually very small numbers of scattered cells and were omitted from analysis.

#### Ipsilateral and contribution input

Data from our initial set of birds confirmed that, like other pallial connections in the bird brain (Reiner et al., 2005), pallial inputs into HF were exclusively ipsilateral. In subsequent birds, we therefore occasionally used the same wavelength of the fluorophore on both hemispheres (7 chickadees). These animals were used only for quantifying pallial inputs into HF.

To compare the strength of input from the ipsilateral and contralateral hemispheres, we computed the following lateralization score (*s*_lat_) for each input brain region:

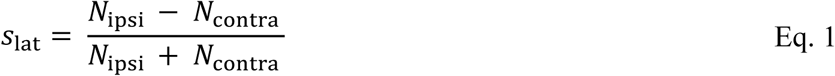

Where *N*_ipsi_ is the number of retrogradely labeled cells in the ipsilateral hemisphere, and *N*_contra_ is the number in the contralateral hemisphere. For every brain region, we calculated the average *s*_lat_ across all CTB injections, regardless of their location in HF.

#### Linear mixed-effects models

We used a linear mixed-effects model to examine how the number of HF-projecting cells in each brain region varied with the coordinate of the injection in HF. This model was fit separately for every input region. Models were fit to the base-10 logarithm of the cell counts, with 1 added to all counts to prevent taking a logarithm of 0:

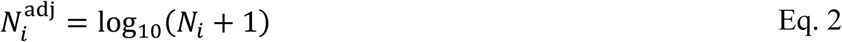

where *N*_*i*_ is the number of labeled cells and *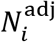* is the adjusted cell count following injection *i*.Here, *i* indexes across all injections across all birds, for which the input region was annotated.

We considered two fixed effects in the model. The first effect was denoted using variable *T* (for transverse axis) and specified whether the injection was into Hp (*T*_*i*_=0) or into DL (*T*_*i*_=1). The second effect was denoted using variable *L* (for long axis) and specified the AP location of the injection in mm. For Hp injections, *L*_*i*_ had a value of -1, 1, 3, or 5 (locations 1-4, Table 1).

For DL injections (locations 5-7, Table 1), *L*_*i*_ was assigned values of 1, 3, and 5, respectively, based on the topographically matched Hp location.

In addition to the two fixed effects, we also considered two random effects. The first random effect was the identity of the bird. This was used to correct for any uncontrolled factors that could affect the count of cells across birds, such as age or the quality of the perfusion. This random effect was assigned coefficients *r*_*1*_, *r*_*2*_, …, *r*_*21*_ for each of the 21 birds used in the experiment. The second random effect was the wavelength of fluorophore used for the injection. This was modeled since the different fluorophores had different signal-to-noise ratios of fluorescence against the surrounding tissue, which could have resulted in systematic differences in cell counts. This random effect was assigned coefficients *s*_488_, *s*_555_, and *s*_647_ for the three different wavelengths of the fluorophore that we used.

For each injection *i*, variable *b*_*i*_ specified the identity of the bird (1 ≤ *i* ≤ 21), and variable *w*_*i*_ specified the wavelength used (488, 555, or 647). Cell counts were then modeled as

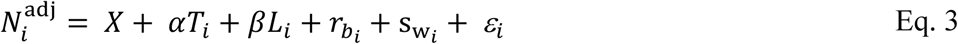

Here, *X* is a fixed offset, *ε*_*i*_ is the observation error for injection *i, α* is the coefficient modeling the effect of the transverse axis location (Hp or DL), and *β* is the coefficient modeling the effect of the long axis coordinate. We used the MATLAB function *fitlme* to fit the coefficients, measure their confidence intervals, and determine whether they significantly differed from zero. These coefficients and p-values are reported in figures 8E and 9E, respectively.

Next, for each input brain region we asked whether there was an axis along which cells preferentially projected to different parts of the HF long axis. Here, we only considered brain regions that had projections to at least three of our four locations along the long axis. This excluded PMI and TnA.

First, we looked for topographic organization of input along the three stereotaxic axes, using a linear mixed-effects model. For this analysis, we only considered injections that labeled at least one cell in the brain region being analyzed. We fit the following linear mixed-effects models to predict how the mean AP coordinate (*A*_*i*_), mean ML coordinate (*M*_*i*_), and mean DV coordinate (*D*_*i*_) varied between injections into different parts of the HF long axis.

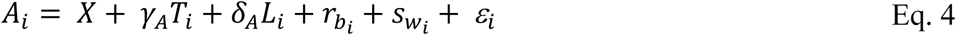

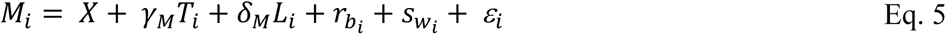

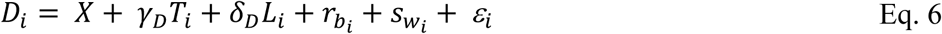

Here, coefficients *γ*_*A*_, *γ*_*M*_, *γ*_*D*_ model the effect of the transverse axis, which we did not analyze. Coefficients *δ*_*A*_, *δ*_*M*_, *δ*_*D*_ model the effect of the long axis. In other words, the vector *v* = (*δ*_*A*_, *δ*_*M*_, *δ*_*D*_) is the direction of the topographic gradient connecting the brain region to the HF long axis.

Next, we projected the location of each labeled cell onto this vector *v*. For each injection *i*, we referred to the mean projected coordinate as *V*_*i*_. This allowed us to compute the center of each retrogradely labeled region along the topographic axis. We used a fourth model for this coordinate

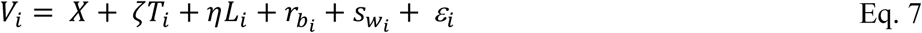

Here, *ζ* models the effect of the transverse axis, which we did not analyze, and *η* models the effect of the long axis. In other words, *η* is the change in the coordinate along the topographic axis of the brain region, for every mm shift in injection coordinate along the HF long axis. We normalized *η* by the overall extent of the brain region along the topographic axis. This extent was estimated using a bird with retrograde labeling from locations 1 and 3, representing almost the entire extent of the structure. Following this normalization, *η* could be expressed in percent change for every mm shift along the HF long axis.

Finally, we noted that the direction of the topographic vector *v* is arbitrary: it is possible to obtain the same model fit by negating both *v* and *η*. We therefore multiplied *η* by -1 if the dot product of *v* with the AP axis (1,0,0) was <0. In other words, *η* was positive if the more anterior parts of the region projected to the more anterior parts of HF.

## Results

### Organization of the avian HF

HF in chickadees consists of Hp, DL, and CDL (Figure 1A). Hp is located in the dorsomedial telencephalon and is clearly defined by a lower cell density compared to the surrounding tissue. DL is situated directly lateral to Hp and has been delineated in our previous work by retrograde tracing from Hp (Applegate et al., 2023). We decided to consider inputs into Hp and DL, since these regions appear to be analogous to the mammalian hippocampus and the entorhinal cortex, respectively. CDL is situated further laterally and is a thin layer extending onto the posterior and lateral surfaces of the brain. In chickadees most of this structure is exceedingly small (often 1-2 cell bodies thick) and was not considered in this study.

**Figure 1:**
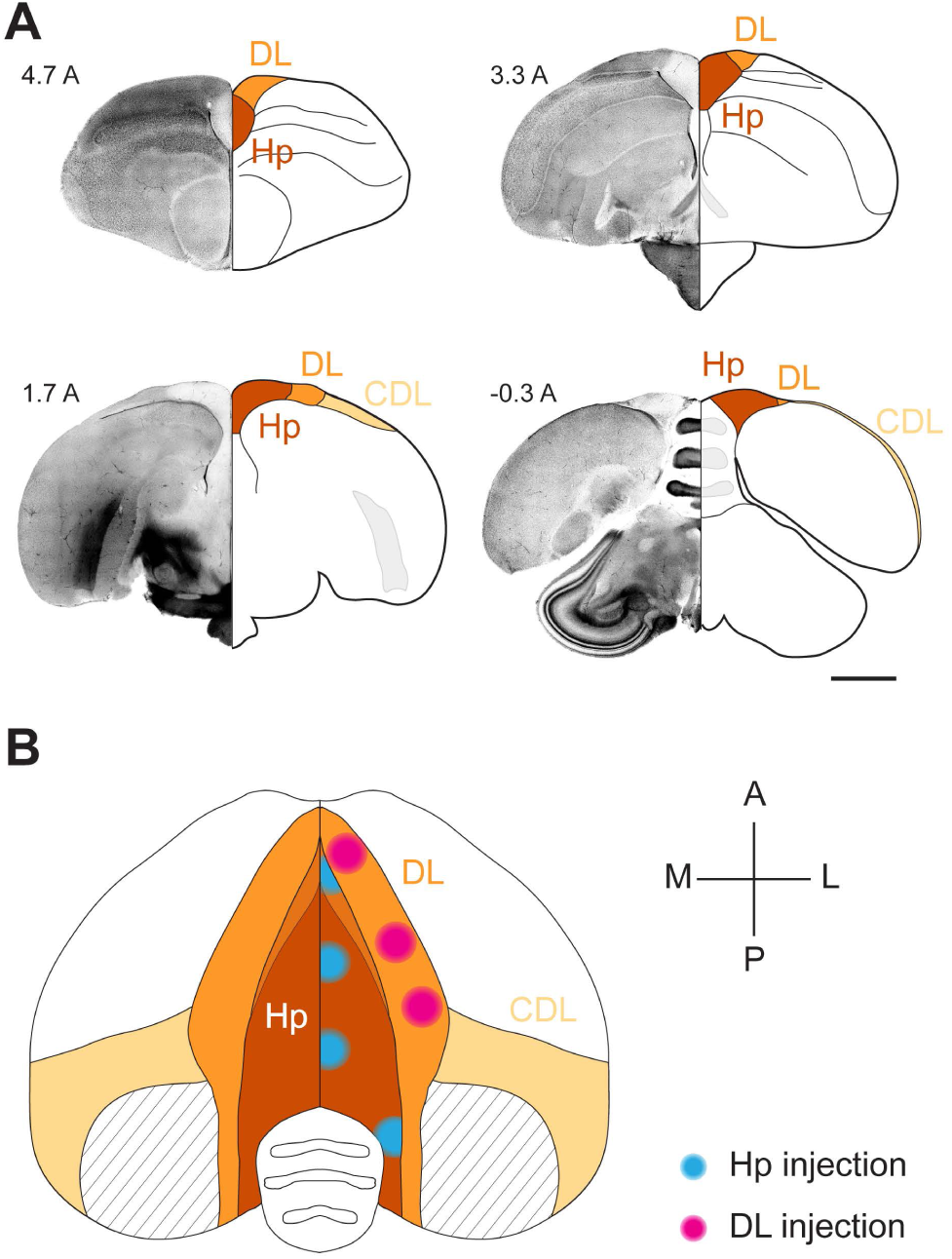
The HF of the black-capped chickadee. (**A)** Coronal sections of the chickadee brain illustrating the boundaries of HF. For each coronal section, the left hemisphere is shown in DAPI stain and the right hemisphere is shown as a section with labeled landmarks. Numeric label indicates the coordinate in mm anterior to lambda. Note the change in cell density between Hp and DL. Also note that, in the anterior-most section, Hp does not extend to the dorsal surface of the brain and is bordered dorsally by DL. In the posterior-most sections, DL is very small, and CDL extends into an extremely thin layer at the brain surface. Scale bar: 2 mm. (**B)** Horizontal projection of the brain indicating the seven CTB injection locations used in this study. Stereotaxic coordinates of each injection are indicated in Table 1. CTB spread (∼750 μm diameter bolus) is indicated by cyan (Hp) and magenta (DL) circles. We did not observe CTB spreading across the midline or across the boundaries of Hp and DL, presumably due to cytoarchitectural differences with the surrounding structures. Striped area indicates the portion of CDL that is a thin fiber tract containing no cell bodies. At anterior positions, DL is dorsal to the hippocampus and overlaps with it in horizontal projection (blended color). At posterior positions, Hp is party dorsal to the cerebellum; the cerebellum is partially occluded at its lateral-most positions.

We used the retrograde tracer cholera toxin subunit B (CTB) to map inputs into HF. We selected injection coordinates that included both Hp and DL, and that spanned most of the AP extent of these structures. For Hp injections, we chose four coordinates at -1A, 1A, 3A, and 5A (mm anterior to lambda, Figure 1B, Table 1). As previously reported, these injections labeled different portions of DL, with anterior DL most strongly innervating the anterior Hp (Applegate et al., 2023). For DL, we therefore chose locations that were topographically matched to the Hp injections (Figure 1B, Table 1). Hp coordinates 1A, 3A, and 5A topographically corresponded to DL coordinates at 2A, 3.5A, and 5.5A, respectively. Injections into the fourth coordinate at -1A labeled an extremely thin posterior tail of DL. This tail was too small for reliable CTB injections and was omitted from the study.

For each of the resulting seven locations (four in Hp and three in DL), we made injections in at least three chickadees (Table 1). In some birds, injections into multiple coordinates were made in different wavelengths of CTB. For every injection, we identified all retrogradely labeled cells using a blob detection algorithm. These cells were classified into different input regions using atlases of other avian species (Karten and Hodos, 1967; Stokes et al., 1974; Puelles et al., 2019). In the next three sections, we will examine all inputs, regardless of which of the seven coordinates they innervated. Afterwards, we will examine the topographic patterning of these inputs to different locations.

### Pallial inputs to HF

We first analyzed inputs to HF from the pallium. These inputs consisted of four distinct nuclei, some of which were large and irregularly shaped. We visualized these inputs in 3D (Figure 2). The most dorsal of these structures was in the hyperpallium, while the other three were in the nidopallium.

**Figure 2:**
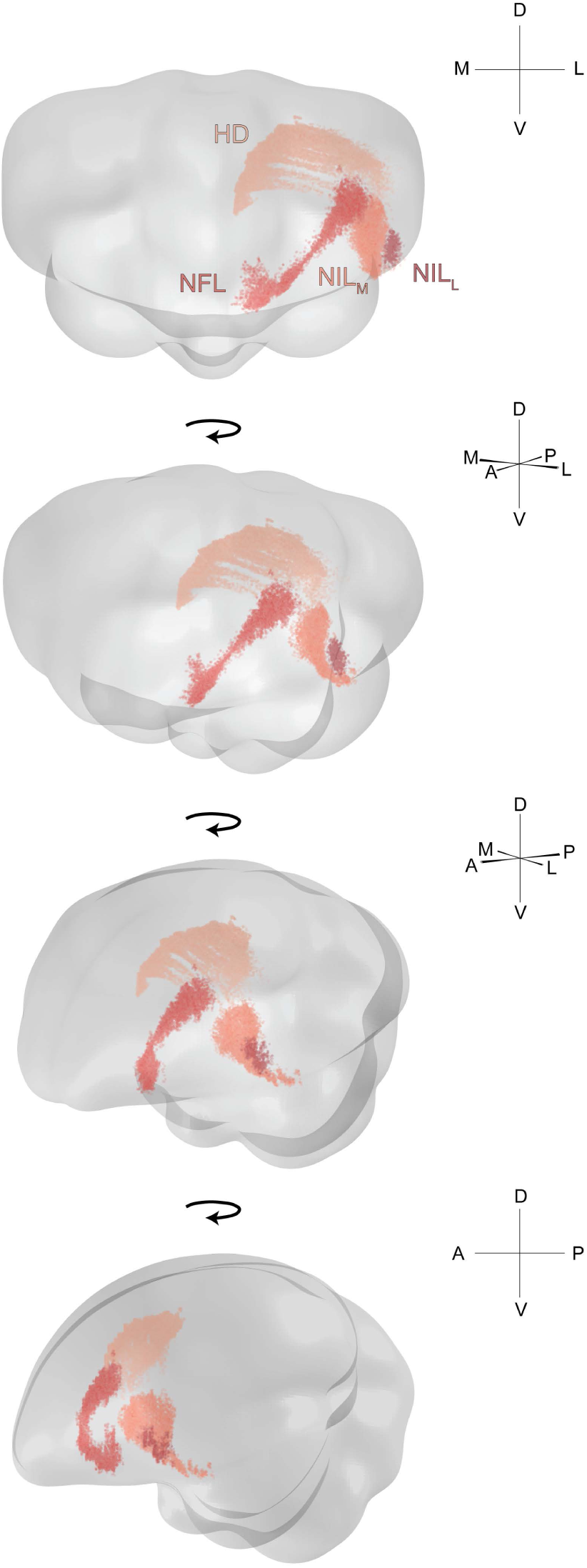
Three-dimensional structure of pallial inputs. Three-dimensional renderings of the chickadee brain, showing pallial regions projecting to HF. First view is from the anterior direction, last view is from the left, and the two intermediate views are rotations by 30° around the DV axis. In all cases the brain is vertically tilted toward the viewer by 10°. A random jitter between -50 µm and +50 µm was added to all AP coordinate for visualization only, since the actual coordinate is discretized by the 100 µm thickness of brain sections. The HD label pools all cells that were retrogradely labeled by three injections in one bird, at locations 2, 3, and 4 (Table 1). The three nidopallial regions are from a separate bird with injections at locations 4 and 7 (Table 1).

The hyperpallial nucleus was HD (Figure 3A). HD is a portion of the visual Wulst constrained dorso-ventrally by two lamina: LFM and LFS (Atoji et al., 2017). In chickadees, HD extended medially to within a few hundred microns of the ventricle and the lateral surface of the brain. It extended roughly 2.5 mm AP. Some prior work has subdivided HD into a medial portion called HD and a lateral portion called HL (lateral hyperpallium) (Atoji et al., 2002; Atoji and Wild, 2005). However, we observed no cytoarchitectural or projection differences along this ML direction, so for the sake of all analyses we classified these cells as HD.

**Figure 3:**
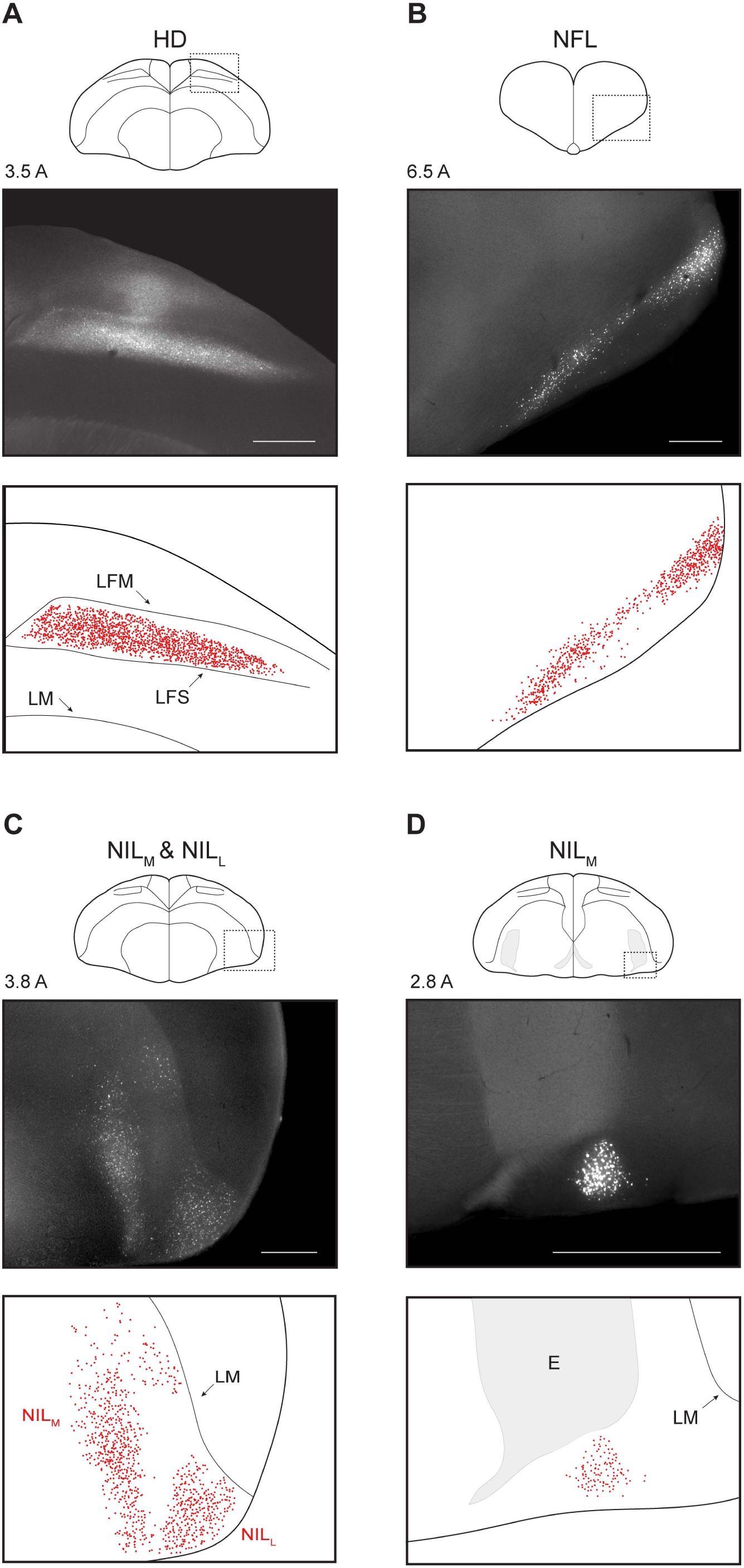
Pallial inputs. (**A-D**) Pallial inputs into HF, as described in the text. For each panel, top plot shows a schematic of a coronal section with the region of interest indicated by a dotted rectangle. Middle plot shows retrograde labeling within the region of interest. Numeric label indicates coordinate of the section in mm anterior to lambda. Bottom plot shows detected neurons and annotated landmarks in the same region of interest. Injections in the four panels were at locations 5, 4, 7, and 4, respectively (Table 1). Scale bars: 500 μm.

The nidopallial nuclei were all in the lateral part of the structure (NL). NL is typically subdivided into frontal, intermediate, and caudal portions based on position relative to the entopallium. The first nucleus we observed was in the frontal potion of NL (NFL, Figure 3B). This extremely anterior population occupied the ventro-lateral edge of the brain. The other two nuclei were in the intermediate portion of NL (NIL) and occupied different ML positions (Figure 3C). We therefore called these nuclei NIL_L_ and NIL_M_. NIL_L_ was bordered ventrally and laterally by the brain surface and dorsally by LM. NIL_M_ was situated more medially, and in its anterior portion spanned the entire extent from LM to the ventral brain surface. In its posterior-most extent, it was constrained dorsally by the entopallium and ventrally by the surface of the brain (Figure 3D). Here, it formed a tight nucleus spanning only a few hundred microns ML and DV.

These pallial nuclei have been described to various extent in the literature. HD is a region of the thalamofugal visual pathway and has been observed as an input to HF in other species (Casini et al., 1986; Atoji et al., 2002; Kahn et al., 2003; Atoji and Wild, 2004; Herold et al., 2019). It is involved in visual memory (Shimizu and Hodos, 1989; Watanabe, 2003; Maekawa et al., 2006) and receives input from other portions of the visual Wulst and from the visual thalamic areas (Medina and Reiner, 2000; Atoji et al., 2017). NL has also been implicated in visual processing as part of the tectofugal pathway (Alpár and Tömböl, 1998; Watanabe, 2003), but may also participate in processing other sensory modalities (Brauth et al., 1987; Brauth and McHale, 1988; Hall et al., 1993). Although NFL has been detected as an input to Hp (Casini et al., 1986; Atoji et al., 2002), neither of the NIL nuclei have been described as inputs to HF in other species. However, it is possible that they overlap with the temporo-parieto-occipital area (TPO), which has been described as an input into to DL (Behroozi et al., 2017).

### Subpallial inputs to HF

We next examined the subpallial inputs to HF. The first two of these were portions of the septum that are known as inputs to HF in other avian species (Casini et al., 1986; Atoji et al., 2002). The anterior-most septal population was NDB (Figure 4A). This nucleus extended about 1 mm AP. In its medial portion, NDB was confined within the fiber tract FDB. At its posterior extent, NDB extended laterally beyond FDB, but remained at roughly the same ventral position. In the posterior-dorsal part of the septum, we also observed a population of labeled cells in MS (Figure 4B). These cells were usually only sparsely labelled, as has been observed in pigeons (Atoji et al., 2002; Atoji and Wild, 2004; Montagnese et al., 2004). In mammals, the septum provides cholinergic and GABAergic input to HF and has multiple functions in HF processing (Stewart and Fox, 1990; Kaifosh et al., 2013; Petersen and Buzsáki, 2020; Schlesiger et al., 2021). Less is known in birds, although the septum in pigeons does play a role in spatial memory (Peterson and Bingman, 2011).

**Figure 4:**
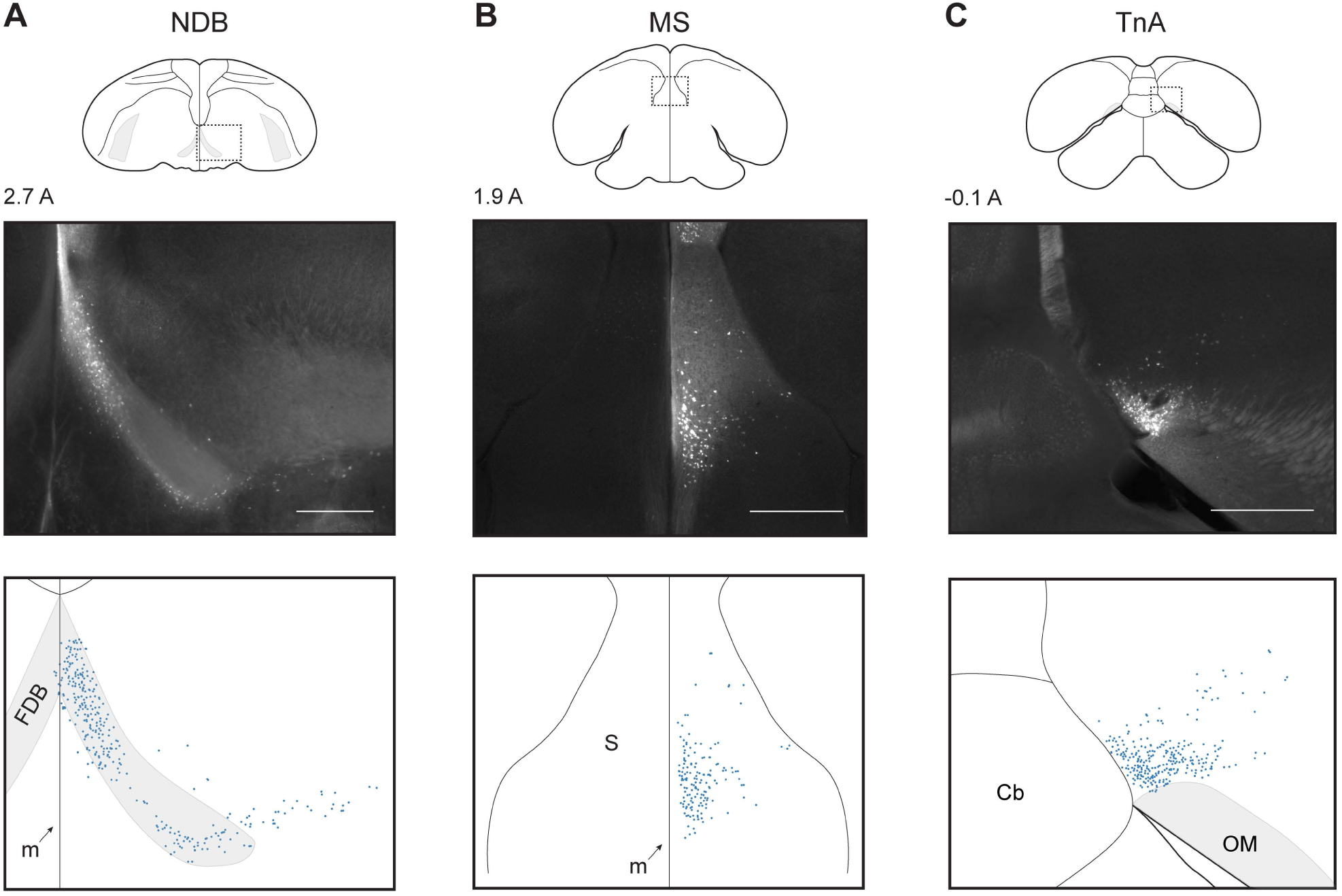
Subpallial inputs. (**A-D**) Subpallial inputs into HF, plotted as in Figure 3. Injections were at locations 3, 2, and 1, respectively (Table 1). Scale bars: 500 μm.

The last subpallial region was TnA in the amygdala (Figure 4C). TnA, also sometimes called the pallial medial amygdalar nucleus (Abellán et al., 2009), was located in the posterior medio-ventral telencephalon, immediately dorsal to OM. In prior avian studies, TnA has been implicated in a variety of social and emotional behaviors, from social feeding to abnormal fear responses (Cheng et al., 1999). We observed labelled cells distributed over 0.5 mm AP. Note that while in other species TnA appears more laterally, its position relative to OM is consistent with our observations (Leutgeb et al., 1996; Cheng et al., 1999).

### Non-telencephalic inputs to HF

We next examined the diencephalic input to HF. In the thalamus we observed four nuclei, all of which have been previously described as inputs to HF (Casini et al., 1986). Going from anterior to posterior, the first of these was SPC (Figure 5A). It was the largest thalamic nucleus, forming a long thin (∼100 μm) sheet that stretched from 1.7A to 0.5A along the DTB. The second and third nuclei were DLM and PMI, respectively (Figure 5B-C). Both extended about 0.5 mm AP and were about 1 mm across in cross-section. The most posterior thalamic nucleus was SRt (Figure 5D). It was bordered laterally by the medial extent of OM and was also the most ventral thalamic population we observed. SRt extended AP for about 300 μm, and was elongated in the ML direction (0.5 mm by 1 mm in cross-section).

**Figure 5:**
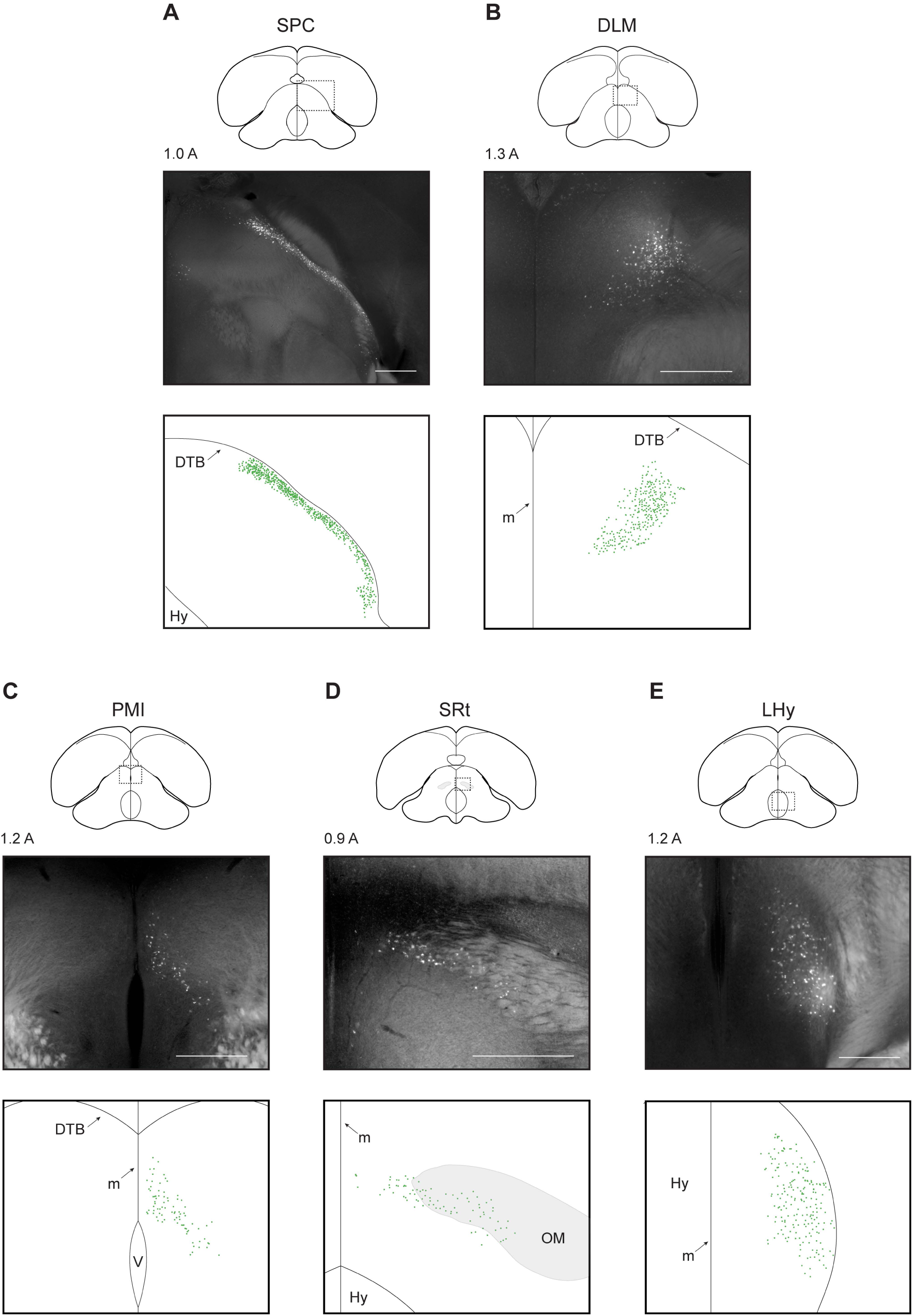
Diencephalic inputs. (**A-E**) Diencephalic inputs into HF, plotted as in Figure 3. Injections were at locations 7, 6, 4, 4, and 4, respectively (Table 1). Scale bars: 500 μm.

The role of these thalamic regions is still largely unknown. SPC, DLM, and SRt all receive visual and somatosensory input, while PMI is considered an auditory structure (Bagnoli and Burkhalter, 1983; Martin Wild et al., 1993; Atoji and Wild, 2006; Miceli et al., 2006; Atoji et al., 2017). A region called DLM is part of the song system (Bottjer et al., 1989), but it is unclear if that structure is the same as the HF-projecting population we labeled.

In the hypothalamus, we observed a lateral population of labeled cells (LHy, Figure 5E). These cells were constrained to the dorso-lateral hypothalamus, and extended for about 800 μm AP. Functionally LHy in both birds and mammals has been implicated in feeding and food-hoarding behaviors (Herberg and Blundell, 1967; Kuenzel, 1972). LHy has also been described as an input to HF in other species (Casini et al., 1986; Atoji et al., 2002).

Next, we observed one midbrain population in the VTA (Figure 6A). This population extended for about 800 μm AP, was bounded ventrally by the brain surface and medially by the midline. As in other species, VTA appeared near the oculomotor nerve (NIII) in coronal sections (Reiner et al., 2004). Functionally, VTA has been implicated in many studies across both birds and mammals as a dopaminergic region involved in reward signaling (Schultz et al., 1997; Gadagkar et al., 2016).

**Figure 6:**
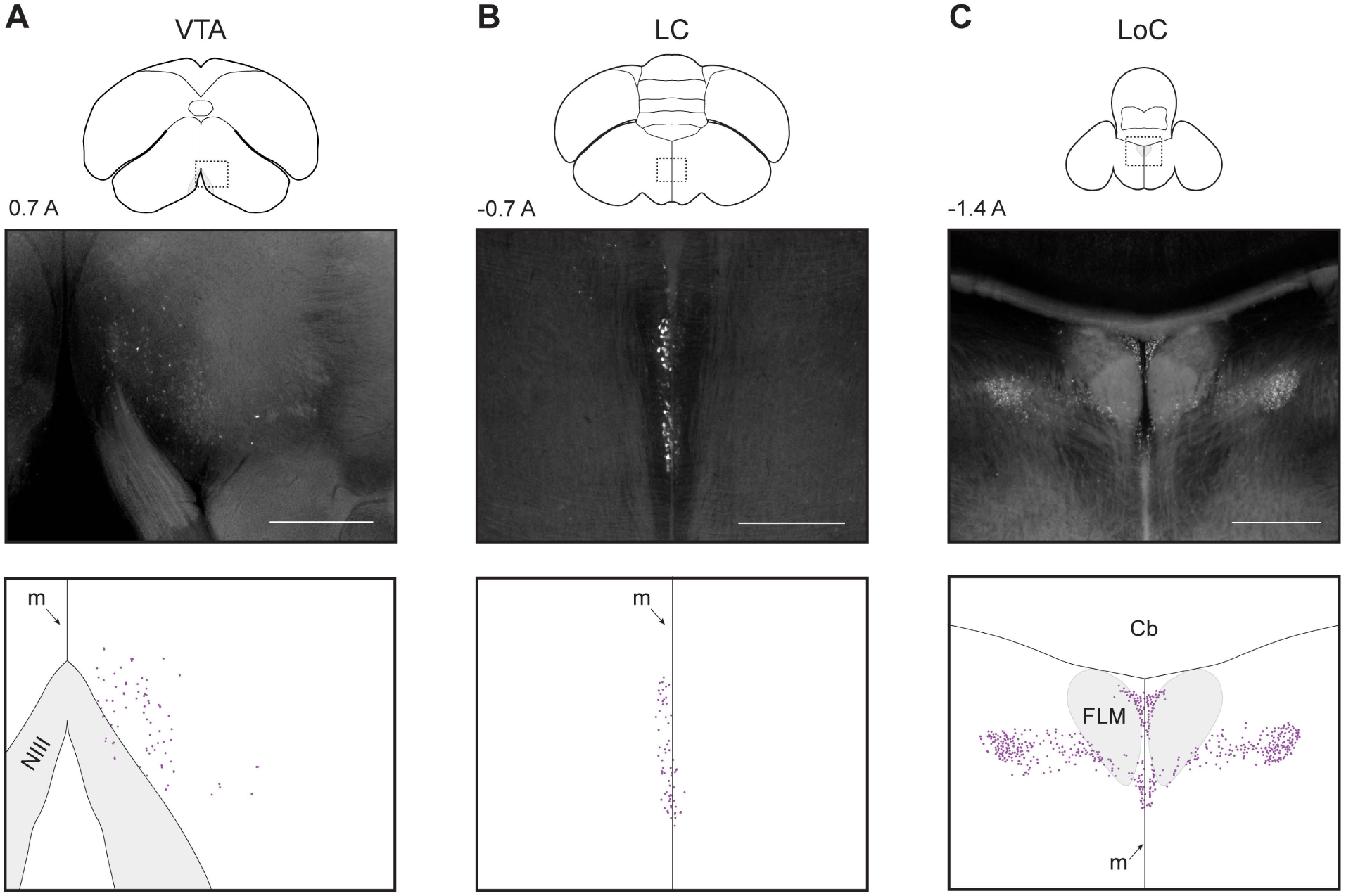
Brainstem inputs. (**A-C**) Brainstem inputs into HF, plotted as in Figure 3. Injections were at locations 4, 1, and 7, respectively (Table 1). Scale bars: 500 μm.

Finally, we observed two populations in the hindbrain, both of which have been previously described as inputs to HF (Casini et al., 1986; Atoji et al., 2002). The more anterior of these was LC (Figure 6B). This nucleus formed a thin sheet (<100 μm) on the midline that extended roughly 700 μm AP and 1 mm DV. This area forms a part of the medial raphe nucleus and has been described as a serotonergic input to HF (Casini et al., 1986; Krebs et al., 1991). The second population was LoC (Figure 6C), a noradrenergic center of the brain (Erichsen et al., 1991; Mello et al., 1998). This input was extremely posterior and extended roughly 700 μm AP. LoC occupied the dorsal portion of the brainstem and surrounded the FLM fiber bundle. At the anterior portion of LoC, cells were closer to the midline (within ∼500 μm) while at the posterior extent, cells were distributed up to 1 mm from the midline.

### Ipsilateral and contralateral contribution of inputs

In the preceding sections, we cataloged the inputs to the chickadee HF. These consisted of four pallial inputs (HD, NFL, NIL_M_, NIL_L_), three subpallial inputs (MS, NDB, TnA), five diencephalic inputs (SPC, DLM, PMI, SRt, LHy), and three brainstem inputs (VTA, LC, LoC). In the mammalian hippocampal circuit, the majority of input is from the ipsilateral hemisphere, though there are some contralateral projections (Goldowitz et al., 1975; Swanson and Cowan, 1977; Burwell and Amaral, 1998). Birds lack a corpus callosum, and in other avian circuits there is even less connection with the contralateral hemisphere than in mammals (Letzner et al., 2016; Wittek et al., 2021). This distinction is meaningful, likely accounting for some extreme lateralization of sensory and motor brain functions in birds (Graves and Goodale, 1977; Clayton, 1993; Vu et al., 1998; Long and Fee, 2008; Martinho et al., 2015; Martinho and Kacelnik, 2016). We were therefore motivated to quantify the relative ipsilateral and contralateral contribution of inputs into HF.

We found striking differences in contralateral input to HF from different brain regions. Some regions, like DLM, projected almost exclusively to the ipsilateral HF (Figure 7A-B). Other regions, like LoC, had similar numbers of neurons projecting to the ipsilateral and contralateral hemispheres (Figure 7C-D). To quantify these differences, we calculated a lateralization score (*s*_lat_) for each input area that ranged from -1 for a completely contralateral input to +1 for a completely ipsilateral input (Figure 7E).

**Figure 7:**
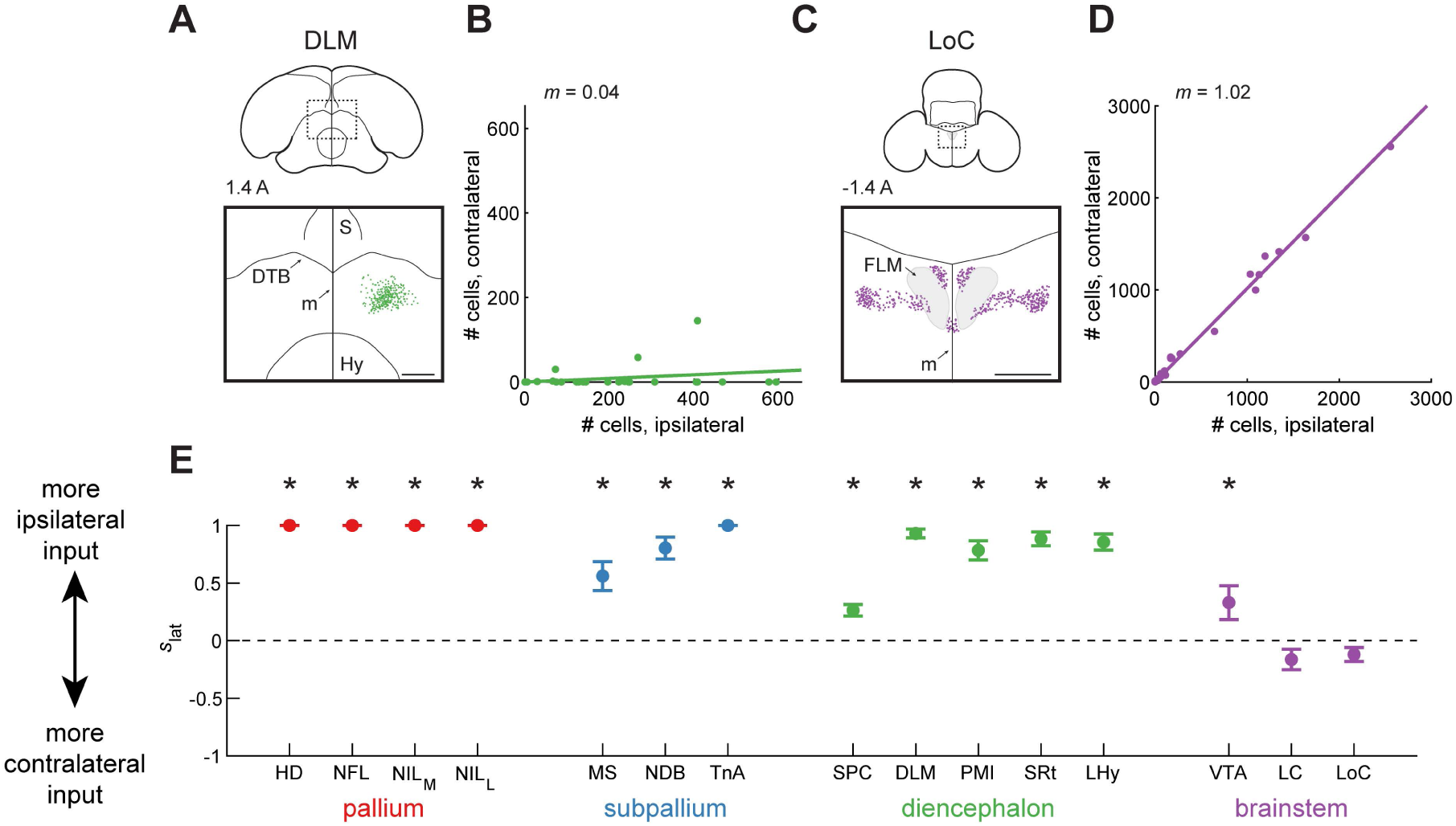
Ipsilateral and contralateral contribution of different inputs to HF. (**A)** Top: schematic of a coronal section, with the region of interest shown by a dotted rectangle. The region of interest is centered on the midline to compare any ipsilateral and contralateral label. Bottom: retrogradely labeled cells. HF receives input predominantly from ipsilateral DLM. Injection was at location 7 (Table 1). Coordinate 1.4 A is in mm anterior to lambda. Scale bar: 500 μm. (**B)** Comparison of the number of cells labeled ipsilaterally and contralaterally. Line is a linear fit with the slope (*m*) indicated. In the case of DLM, slope is smaller than 1, indicating that there was significantly more ipsilateral label (p<0.001, Wilcoxon signed-rank test). (**C)** Labeling of LoC, shown as in (A). HF receives input from both contralateral and ipsilateral hemispheres. Injection was at location 7 (Table 1). Scale bar: 500 μm. (**D)** Analysis LoC, shown as in (B). The slope is not significantly different from 1, indicating similar labeling of ipsilateral or contralateral cells (p=0.14, Wilcoxon signed-rank test). (**E)** Ipsilateral score *s*_lat_, defined as 1 for a completely ipsilaterally projecting region and -1 for a completely contralaterally projecting region. Mean ± SEM is shown for all regions. Asterisks indicate regions for which *s*_lat_ was significantly different from 0 (p<0.05). Pallial, subpallial, diencephalic, and midbrain (VTA) inputs were predominantly ipsilaterally projecting, while the hindbrain (LC and LoC) contributed input from both the ipsilateral and the contralateral hemispheres.

These scores had interesting patterns for regions from the same part of the brain (Figure 7E). Like in most avian circuits, there was no contralateral input from any of the pallial regions (Nottebohm et al., 1976; Schmidt et al., 2004). In contrast, subpallial and diencephalic areas provided some contralateral input, though they also projected primarily ipsilaterally. The most notable contralateral projections among these areas were from MS and SPC. The brainstem regions had the most balanced projections to the two hemispheres. In particular, neither of the hindbrain inputs (LC and LoC) showed significant differences between contralateral and ipsilateral labeling. Collectively, our results indicate a hippocampal circuit that has only a few regions for cross-hemispheric routing of information.

### Topography along the transverse axis of HF

We next asked how projections from these input areas differed between Hp and DL. Our prior work has already described some differences in the relative strength of pallial inputs to these structures (Applegate et al., 2023). Here, we provide a more detailed analysis of the different pallial subregions, and we also analyze all of the non-pallial inputs.

We observed differences in projections to DL and to Hp. To quantify these observations, we applied a linear mixed-effects model (Gałecki and Burzykowski, 2013), which measured the difference in the number of cells labeled by DL and Hp injections after compensating for other factors that could affect these numbers. The factors we considered were the AP position of the injection, the wavelength of the fluorophore, and the identity of the bird. Note that the number of labeled cells is not a perfect measure of the connection strength, since it can be influenced by factors like fibers of passage or the efficacy of CTB take-up.

Some inputs preferentially innervated DL. For example, only the ventral portion of HD projected to Hp, while the entire DV extent of HD projected to DL (Figure 8A). This difference led to nearly a 3-fold increase in the number of labeled cells (p<0.001, linear mixed-effects model, Figure 8B). In some other brain regions, there were more direct inputs to Hp than to DL. For example, LHy had a markedly weaker projection to DL, with several injections labeling no cells (Figure 8C-D, p<0.01).

**Figure 8:**
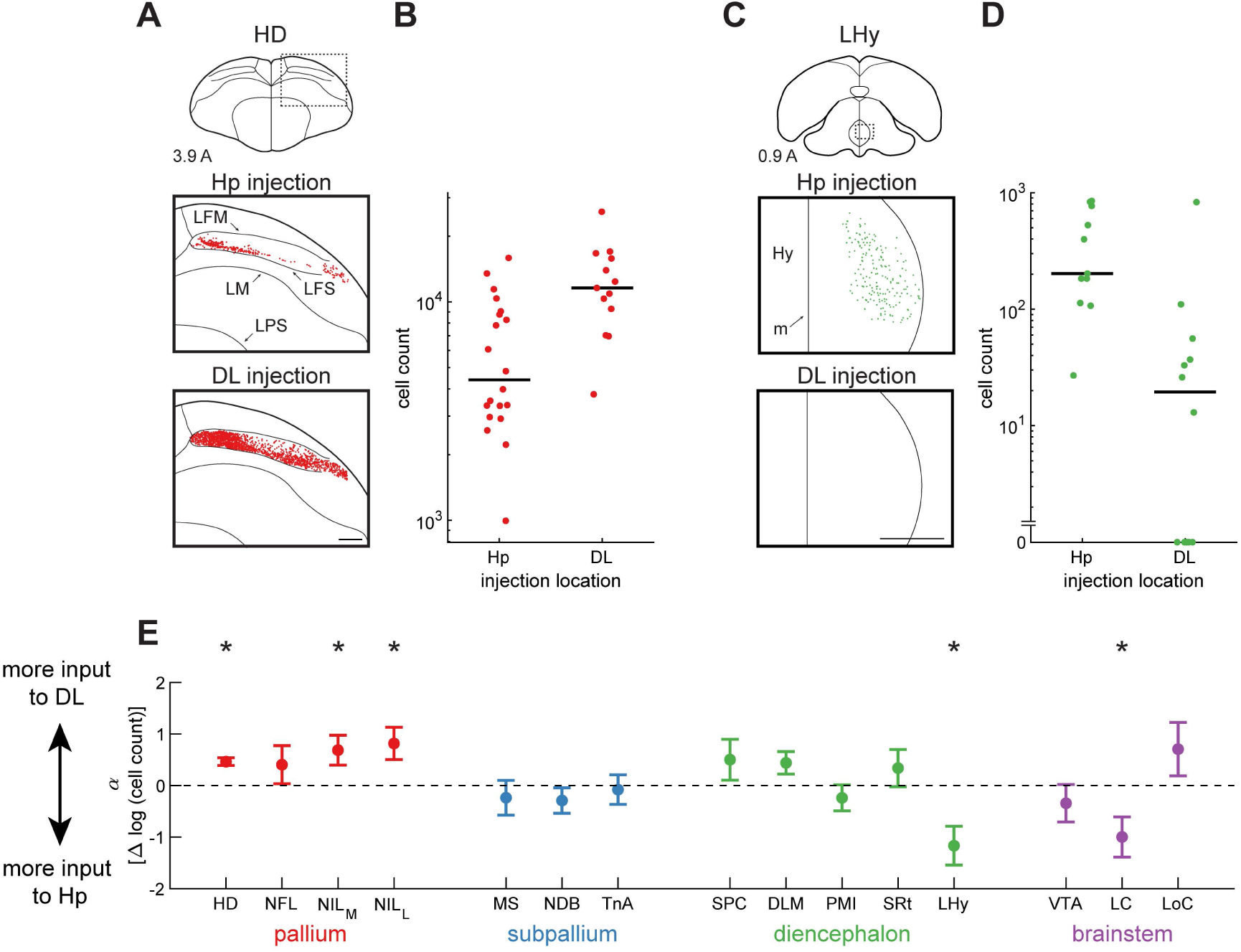
Inputs to HF differentially target DL and Hp. (**A** Top: schematic of a coronal section, with the region of interest shown by a dotted rectangle. Middle and bottom: retrogradely labeled neurons from a paired Hp injection and DL injection. Examples are from the same coronal section of the same brain. There were more cells labeled by the DL injection than by the Hp injection. Injections were at locations 3 and 6 (Table 1). Coordinate 3.9 A is in mm anterior to lambda. Scale bar: 500 μm. (**B)** The number of cells retrogradely labeled by each Hp and DL injection. Data are pooled from all positions along the long axis. In the case of HD, there were significantly more HD cells projecting to DL than to Hp (p<0.001, linear mixed-effects model). Horizontal lines indicate medians. (**C)** Labeling of LHy, shown as in (A). Note the absence of any retrogradely labeled cells from the DL injection. Injections were at locations 4 and 7 (Table 1). Scale bar: 500 μm. (**D)** Analysis of LHy, shown as in (B). There were significantly more cells projecting to Hp than to DL (p<0.01, linear mixed-effects model). (**E)** Parameter *α* of the linear mixed-effects model, which quantifies the difference in retrograde labeling between DL and Hp injections. An increase of 1 in this parameter corresponds to a 10-fold increase in the number of cells retrogradely labeled by DL injections relative to Hp injections. Mean ± bootstrap SEM is shown for all regions. Asterisks indicate regions for which *α* was significantly different from zero (p<0.05). Three of the four pallial regions projected predominantly to DL while two subcortical regions projected predominantly to Hp.

Across all of the brain regions, the pallial nuclei HD, NIL_M_, and NIL_L_ had stronger projections to DL (Figure 8E). These results are consistent with the mammalian literature, where the entorhinal cortex receives more direct cortical input than the hippocampus (Burwell and Amaral, 1998). Only LHy and LC preferentially innervated Hp (Figure 8E). This result is also consistent with observations in rodents. Rodent LHy primarily sends connections to the subiculum in the hippocampus, and not to the entorhinal cortex (Hahn and Swanson, 2012). The rodent medial raphe nucleus, where LC is located, also innervates rat hippocampus more than the entorhinal cortex (Köhler and Steinbusch, 1982). These results are further evidence of the analogy between DL and the entorhinal cortex, and suggest additional similarities in organization between the mammalian and avian circuits.

### Topography along the long axis of HF

We next asked whether input regions exhibited topography of their connections along the HF long axis. We considered two possible forms of topography. First, an input region could preferentially target the anterior or posterior HF. In this case, anterior and posterior injections would retrogradely label different numbers of cells. Second, different portions of the same input region could project to different sections of the HF long axis. In this case, the coordinates of retrogradely labeled cells would be different between anterior and posterior injections.

We first looked at how the number of labeled cells varied with injection location. Again, we applied a linear mixed-effects model to quantify this effect. In some regions, anterior HF injections resulted in more labeled cells. For example, thalamic input from PMI was exclusively limited to the anterior HF (Figure 9A-B, p<0.001). In contrast, amygdalar input from TnA exclusively targeted the posterior HF (Figure 9C-D, p<0.001). We found that the majority of input regions preferentially targeted either the anterior or the posterior HF (Figure 9E). As before, interesting patterns emerged across divisions of the brain. For example, all pallial and all thalamic areas preferentially innervated the anterior HF. In contrast, two of the subpallial regions, TnA and MS, were the only inputs that preferentially targeted the posterior HF.

**Figure 9:**
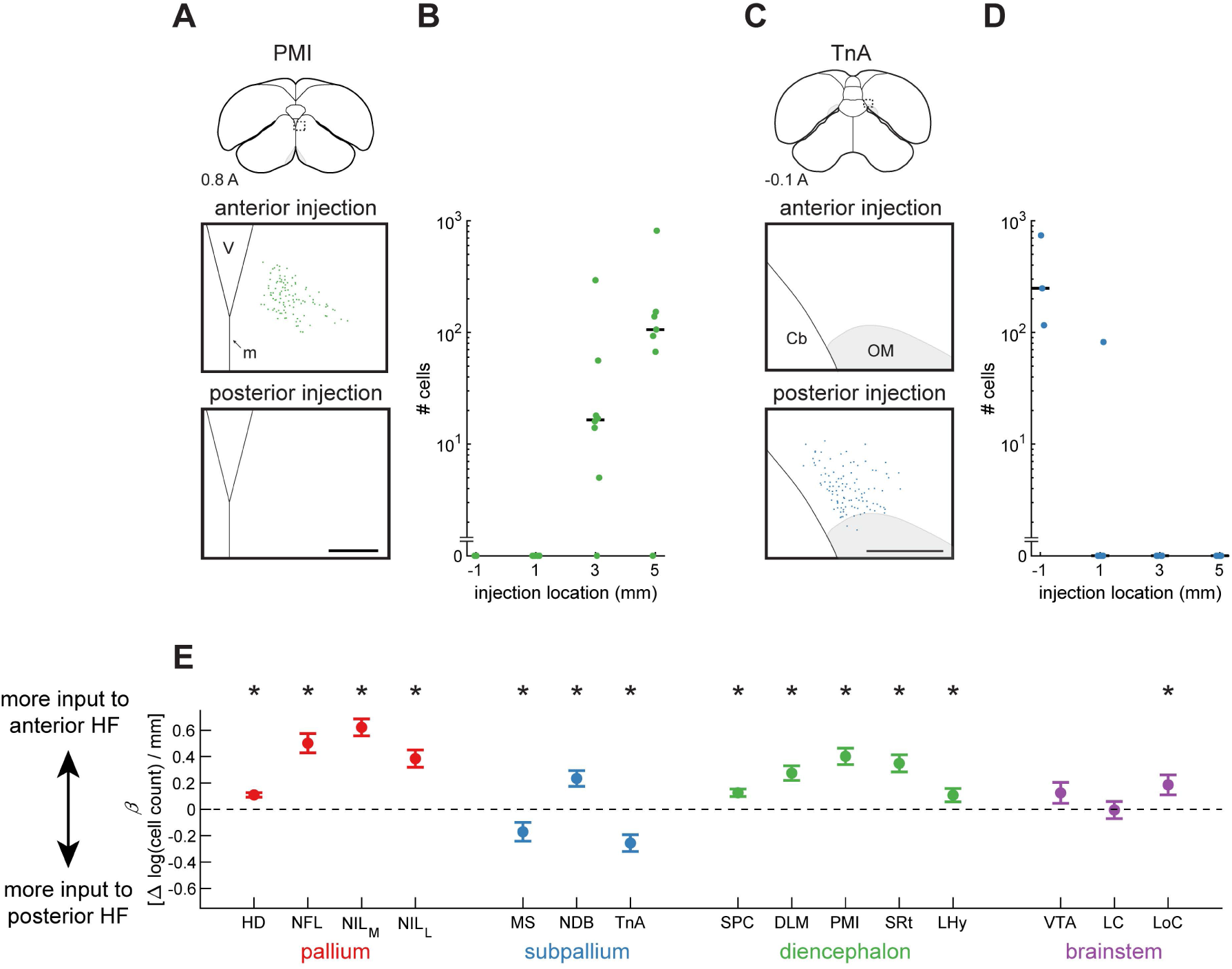
Strength of inputs to HF vary along the long axis. (**A**) Top: schematic of a coronal section, with the region of interest shown by a dotted rectangle. Middle and bottom: retrogradely labeled neurons from paired anterior and posterior injections. Examples are from the same coronal section of the same brain. Labeling was present exclusively from the anterior injections. Injections were at locations 2 and 4 (Table 1). Coordinate 0.8 A is in mm anterior to lambda. Scale bar: 250 μm. (**B**) Number of cells retrogradely labeled by each injection, organized by position along the long axis of HF. Data are pooled from all Hp and DL injections. For Hp points, injection coordinate is in mm anterior to lambda. For DL points, this coordinate indicates the location of the topographically matched Hp location (Table 1). In the case of PMI, there were significantly more cells labeled by the more anterior injections (p<0.001, linear mixed-effects model). Horizontal lines indicate medians. (**C**) Labeling of TnA, shown as in (A). TnA was labeled only by posterior hippocampal injections. Injections were at locations 1 and 4 (Table 1). Scale bar: 250 μm. (**D**) Analysis of TnA, shown as in (B). There were significantly more cells labeled by the more posterior injections (p<0.001, linear mixed-effects model). (**E**) Coefficient *β* of the linear mixed-effects model, which quantifies the effect of injection location on the number of retrogradely labeled cells. An increase of 1 in this parameter corresponds to a 10-fold increase in the number of retrogradely labeled cells for every mm anterior shift of the injection location in HF. Mean ± bootstrap SEM is shown for all regions. Asterisks indicate regions for which *β* was significantly different from zero (p<0.05). With the exception of two brainstem regions, all inputs had a significant increase or decrease in the number of cells projecting to different locations along the long axis.

Finally, we asked whether different portions of these regions projected to different parts of the long axis. We observed some regions with noticeable differences; for example, anterior and posterior HD projected to more anterior and posterior parts of HF, respectively (Figure 10A-B). To quantify this phenomenon, we again used a linear mixed-effects model. We determined the relationship between the location of the HF injections (along the AP axis) and the average stereotaxic coordinate of retrogradely labeled cells. This model quantified the magnitude of this topographic gradient as the amount of shift in the position of labeled cells for every millimeter shift in the injection location. We also quantified the direction of this gradient in stereotaxic coordinates.

**Figure 10:**
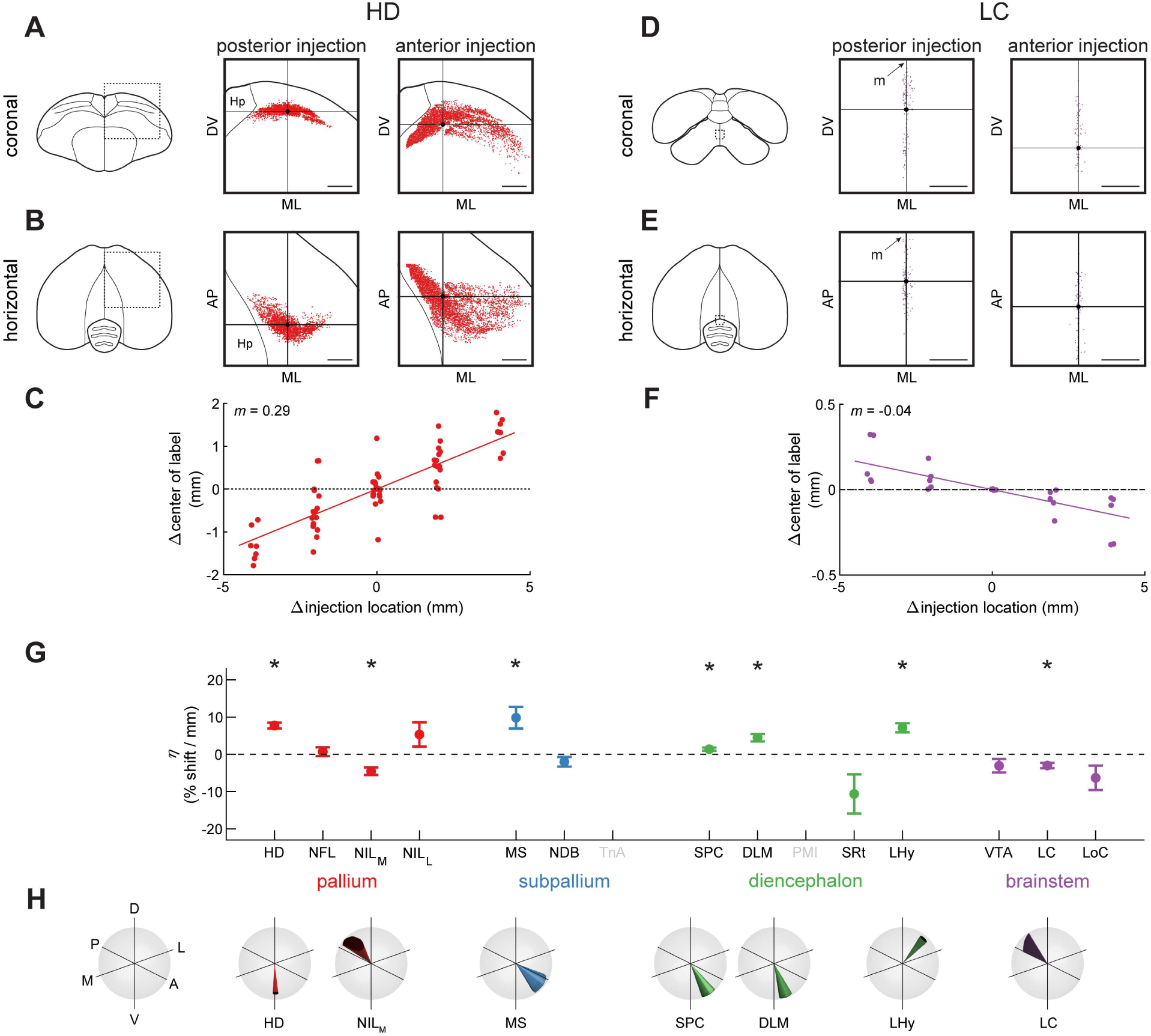
Topographic differences of inputs to HF. **(A)** Left: schematic of a coronal section, with the region of interest shown by a dotted rectangle. Center and right: retrogradely labeled neurons from paired posterior and anterior injections at locations 2 and 4 (Table 1) respectively. This is a projection of all labeled HD cells from all coronal sections of the brain, not only the section shown on the left. Horizontal and vertical lines denote the mean coordinates of the labeled cells, respectively. Scale bars: 1 mm. **(B)** Same as (A), but for the horizontal projection of the brain. A random jitter between -50 µm and +50 µm was added to all AP coordinate for visualization only, since the actual coordinate is discretized by the 100 µm thickness of brain sections. Scale bars: 1 mm. **(C)** Relationship between injection location into HF and the location of retrogradely labeled cells in HD. Every symbol indicates a pair of injections at different coordinates in the same brain. X-axis is the difference in injection coordinate along the AP axis, and y-axis is the difference in the mean coordinate of labeled cells in HD, measured along the main topographic axis. Slope *m* indicates that for every 1 mm shift of the hippocampal injection in the anterior direction, the mean of HD label shifts 0.29 mm along the main topographic axis. Note that the plot is symmetric around (0,0) because for each pair of injections we plot both the comparison of the anterior injection to the posterior one and vice versa. **(D-F)** Analysis of LC, shown as in (A-C). Note that anterior injections labeled cells that were significantly more posterior (p<0.005, linear mixed-effects model). Injections were at locations 2 and 4 (Table 1). Scale bars: 250 μm. **(G)** Coefficient *η* of the linear mixed-effects model, which quantifies the effect of injection location on the location of retrogradely labeled cells. This coefficient is normalized by the overall size of the brain region and expresses percent shift along the topographic axis for every mm anterior shift of the injection location. Asterisks indicate regions for which *η* was significantly different from zero (p<0.05). TnA and PMI were not included in this analysis because in both cases only two of the injection locations labeled any cells. About half of the regions are significantly topographic in this analysis. **(H)** The orientations of the topographic axes for the brain areas that were significantly topographic in (G). Cone indicates the direction of the topographic axis in stereotaxic coordinates. Width of the cone indicates standard error of this estimate (note that standard error can be different in different directions, making the base of the cone an ellipse).

Using this approach, we found that different portions of HD indeed projected to different parts of the HF long axis (Figure 10C, p<0.001) with roughly a 300 μm shift for every mm shift in injection location. Repeating this procedure for other brain regions revealed other interesting topographies. For example, anterior LC preferentially targeted the more posterior HF locations (Figure 10D-F, p<0.005) with roughly a 40 μm shift per mm shift in injection location. To compare topographies across regions, we normalized the magnitude of each topographic gradient by the size of the brain structure along the direction of the gradient. We expressed these coefficients as the percent change in position for every mm change in injection location (Figure 10G-H).

About half of the input regions exhibited a significant topographic gradient. Again, some of these differences suggest interesting parallels to the mammalian circuit. Two of the pallial regions had a topographic gradient with the HF long axis. These types of patterns are common in mammalian cortical regions. For example, the orbitofrontal cortex is connected topographically to the lateral entorhinal cortex (Kondo and Witter, 2014). Some subcortical regions also had mammalian-like organization. As in our results, anterior-dorsal LHy in rodents preferentially innervates the septal hippocampus, while posterior-ventral LHy preferentially innervates the temporal hippocampus (Hahn and Swanson, 2012).

## Discussion

In this study we used retrograde tracing to map inputs into the chickadee HF. By varying our injection location, we examined how inputs differ along both the transverse and the long axes of HF. Input regions were found in several parts of the brain, and were generally similar to those described in other avian species and in mammals. We discovered that many of these inputs were topographic, with some striking similarities to the mammalian hippocampal system. These kinds of similarities are particularly surprising given the major changes in neural architecture between these species, including the lack of a layered cortex or a corpus callosum in birds. In addition to the similarities, we found some notable differences between chickadees and other species. These unique features of the anatomy will be particularly interesting to examine in the future, as they may contribute to the specialized memory functions of food-caching birds.

Our injections spanned almost the entire volume of HF in chickadees. We therefore think that we have identified nearly all inputs into this structure. There are two possible exceptions.

First, we did not inject into the most ventral *“*V*”* portion of Hp (Székely, 1999; Atoji and Wild, 2004, 2006). Previous studies in other species have identified only MS as an extrinsic input into this region (Székely, 1999; Atoji and Wild, 2005). Second, we did not inject into CDL. We previously found that this region is topographically connected to Hp and appears to be similar to the lateral entorhinal cortex (Applegate et al., 2023). In other avian species, it is associated with the olfactory system and receives input from the piriform cortex (Atoji and Wild, 2005). In chickadees, CDL and the rest of the olfactory system are exceedingly small (Bang and Cobb, 1968) and could not be studied with our tracing protocols. This is consistent with chickadee behavior, as they are not known to use olfaction for any memory tasks (Sherry et al., 1981; Herz et al., 1993). However, if CDL is to be considered part of HF, there may be an additional vestigial olfactory input not reported here.

### Lateralization of the hippocampal circuit

The chickadee hippocampal circuit was extremely lateralized. All of the pallial inputs and the majority of subpallial and thalamic inputs in our data were predominantly ipsilateral. This anatomical segregation likely explains lateralized learning in certain memory tasks. Although chickadees have some binocular overlap of their visual field (Moore et al., 2013), they use monocular vision for most perceptual tasks (Martin, 2009). When chickadees cache food with one eye covered by an eye patch, they are unable to find their caches shortly afterwards if the patch is moved to the other eye (Sherry et al., 1981; Clayton, 1993). This indicates that some crucial aspect of cache memory is not shared between hemispheres. Such a pattern of *‘* ipsilateral learning*’* is not unique to chickadees. In other visual tasks, like reversal learning, homing in pigeons, and imprinting in ducklings, there is also a lack of interocular transfer of learned information (Graves and Goodale, 1977; Martinho et al., 2015; Martinho and Kacelnik, 2016).

Interestingly, there is evidence that this transfer does happen after longer periods of time. For example, chickadees in the eye-patch experiment are able to find caches after 24 h (Clayton, 1993). It is unknown how this offline transfer happens. Our analysis has identified some input regions with particularly strong contralateral projections, which could play a role in this process. Another possible mechanism is via the hippocampal commissure (Atoji et al., 2002; Letzner et al., 2016).

### Similarities to mammalian circuitry

Organization of inputs into HF had notable similarities between chickadees and mammals. One of these similarities was in the transverse axis, which separates Hp from DL in birds and from the entorhinal cortex in mammals. As in rodents, we found that hypothalamic inputs preferentially targeted Hp (Hahn and Swanson, 2012). In contrast, pallial inputs preferentially targeted DL, much like they target the entorhinal cortex in mammals (Burwell and Amaral, 1998). Previous work has demonstrated numerous anatomical and physiological similarities of DL and the entorhinal cortex (Atoji and Wild, 2004; Abellán et al., 2014; Applegate et al., 2023). Our findings add further evidence of the equivalence between these regions across species.

In addition, we found similarities in the topography of inputs with the HF long axis. This topography was notable because it might contribute to the known organization of functions along this axis. We found that thalamic and some pallial inputs preferentially innervated the anterior HF in chickadees, much like they innervate the dorsal hippocampus in rodents. As in rodents, some of these inputs are associated with the visual system (Karten et al., 1973; Miceli et al., 2006; Atoji et al., 2017), which is critically important for spatial computations. These inputs may contribute to the selective abundance of spatially tuned cells in the anterior and dorsal hippocampi of chickadees and rodents, respectively. In contrast, we found a highly selective innervation of the posterior HF by the amygdala. This is equivalent to the amygdalar inputs into the rodent ventral hippocampus (Van Groen and Wyss, 1990; Petrovich et al., 2001). The rodent ventral hippocampus is involved in emotional and stress-related behaviors, and a similar function has been proposed for this region in birds (Bouillé and Baylé, 1973; Smulders, 2017). Overall, the topographies we found support the idea that inputs play a major role in the functional organization of the long axis, in both birds and mammals.

How might all these similarities between birds and mammals emerge? One possibility is that both the transverse and the long axes are derived from a common amniote precursor, and that these gradients of input have been preserved over millions of years (Witter et al., 2017; Herold et al., 2019). Indeed, similar transcription factors are expressed along the transverse axes across species (Abellán et al., 2014; Tosches et al., 2018). Conversely, similar topographies could have emerged through convergence. A functionally segmented hippocampus may confer some advantage to the animal. Theoretical work has shown that having related but distinct functional regions can aide in computation (Rolls and Webb, 2014).

### Specialization in a food-caching bird

In addition to the similarities, we also found notable differences between chickadees and other species. In mammals, the difference between inputs to the hippocampus and the entorhinal cortex is dramatic. Nearly thirty cortical regions project to the entorhinal cortex, while only two of them send weak inputs directly to the hippocampus (Dolorfo and Amaral, 1998). In chickadees, the difference between Hp and DL was more subtle. All DL-projecting regions also targeted Hp, albeit often less strongly. This result suggests a less pronounced specialization of HF subregions in birds.

Another striking difference with previous avian studies is the input from the nidopallium. Work in other species has identified a relatively small input into HF from the frontal part of the nidopallium, NFL (Atoji and Wild, 2004). We found a much more expansive input region that also included parts of the NIL and that was highly structured into several distinct nuclei. Parts of NL receive input from the tectrofugal visual pathway and possibly other sensory pathways (Hall et al., 1993; Alpár and Tömböl, 1998; Watanabe, 2003). Other than this, not much is known about any of the regions we identified. Given that these inputs appear to be enhanced in chickadees, it will be interesting to investigate their possible role in food-caching behaviors. The topographic patterns we describe also appear to be partly different from prior work.

Some of our results, such as the topography of NDB and TnA, were previously reported in pigeons (Kraniak and Siegel, 1978; Casini et al., 1986; Atoji et al., 2002; Herold et al., 2019). However, most inputs to the avian HF were previously described without any topography. Most strikingly, some of the clearest topographic patterns we observed, such as the AP gradient of input in HD, have been explicitly described as absent in pigeons (Herold et al., 2019). Some other topographies appear to be stronger in chickadees. For example, TnA has been described as innervating the posterior half or posterior two-thirds of Hp in several studies (Casini et al., 1986; Atoji et al., 2002; Herold et al., 2019). In our data, TnA only innervated the posterior-most quarter of HF. What might be driving these differences between species?

One possibility is that some topographic patterns are harder to observe in other birds. In chickadees, we took advantage of the large hippocampus to study multiple injection coordinates, spaced 2 mm apart. We also quantified the location of every labeled cell and compensated for possible sources of noise like inter-individual variability. Without this quantification, or in an animal with a smaller hippocampus and fewer injection coordinates, it may be hard to detect some of the topographies.

Another possibility is that HF in chickadees is segregated into functional domains more strongly than in other species. Chickadee hippocampus needs to support an immense memory load to remember not only the location of caches, but also their content and other salient features (Cowie et al., 1981; Sherry and Vaccarino, 1989; Clayton and Dickinson, 1998). It is possible that increased computational demands are facilitated by specialized domains similar to those in rodents (Risold and Swanson, 1996; Fanselow and Dong, 2010; Strange et al., 2014). In chickadees some inputs, like PMI and TnA, project to completely non-overlapping parts of HF. It remains to be seen what role some of these extreme topographies play in the hippocampal specialization of food-caching birds.

## Acknowledgements

We thank Stephanie Hale, David Scheck, and Luke Hammond for technical assistance; the Black Rock Forest Consortium, Timothy Green, and Jennifer Scribner, for field site help. Imaging was performed with support from the Zuckerman Institute*’*s Cellular Imaging platform. This work was supported by the Beckman Foundation Young Investigator Award, the New York Stem Cell Foundation — Robertson Neuroscience Investigator Award, NIH Director*’*s New Innovator Award (DP2-AG071918), NIH training grant (T32 EY013933, to MCA), NSF Graduate Research Fellowship Program (to MCA).

